# Influence of the Glymphatic System on α-Synuclein Propagation: Role of Aquaporin-4 and the Dystrophin-Associated Protein Complex

**DOI:** 10.1101/2024.09.11.612272

**Authors:** Douglas M Lopes, Sophie K Llewellyn, Sheila E Bury, Jiazheng Wang, Jack A Wells, Matthew E Gegg, Guglielmo Verona, Mark F Lythgoe, Ian F Harrison

## Abstract

Propagation and aggregation of prion proteins, such as tau and α-synuclein (αSyn), are key pathological features of neurodegenerative diseases. Extracellular clearance pathways, such as the glymphatic system, may play a crucial role in the removal of these toxic proteins from the brain. Primarily active during sleep, this system relies on aquaporin-4 (AQP4) water channel expression and polarisation to astrocytic endfeet, facilitating interstitial solute clearance. Glymphatic dysfunction has recently been implicated in Parkinson’s disease, however the precise mechanisms underlying the pathogenic effect of this dysfunction remain unclear. This includes how impaired glymphatic function influences αSyn propagation dynamics, and the role of propagating αSyn itself on glymphatic function.

In this study, we used a mouse model of αSyn propagation to elucidate the impact of αSyn aggregation on glymphatic function, by measuring CSF-ISF exchange and assessing AQP4 and associated endfoot complex proteins in the brain over time and across different regions. Our results show that direct injection of αSyn pre-formed fibrils leads to reduced expression of the AQP4 endfoot complex, but propagation of endogenous αSyn induces an enhancement of glymphatic function suggesting compensatory upregulation in response to increasing endogenous αSyn load. To determine the influence of glymphatic dysfunction on αSyn propagation dynamics, we then employed a pharmacological approach to inhibit glymphatic function in this model. Acute glymphatic inhibition significantly reduced brain to CSF αSyn clearance, and chronic treatment exacerbated αSyn pathology, neurodegeneration, and motor behavioural deficits in mice. Together our findings show that αSyn clearance and propagation are modulated by glymphatic function and suggest that AQP4 complex dysregulation may contribute to glymphatic impairment associated with Parkinson’s diseases.

**Summary for the non-scientific community:** The glymphatic system clears brain waste during sleep. Lopes et al. show that α-synuclein, a protein linked to Parkinson’s, is cleared by this system. Using a mouse model of the disease, they suggest that aquaporin-4 water channels may impair glymphatic function, contributing to α-synuclein buildup in patients’ brains.

## Introduction

The accumulation of misfolded amyloid-structured proteins is a pathological hallmark of neurodegenerative disorders such as Alzheimer’s disease, Parkinson’s disease, and Huntington’s disease [1]. Although each of the proteins known to accumulate in these disorders are molecularly distinct, they share a common ‘prion-like’ property, giving them the ability to ‘seed’ pathology and spread from cell-to-cell across the neuronal circuit, contributing to the stereotypic spread and progressive nature of pathology in these disorders [2–4]. The mechanisms which govern the spread of intracellular hallmark proteins are still being understood [3, 5–9], but evidence is mounting to suggest that extracellular clearance pathways might represent therapeutic potential for management of these toxic proteins, limiting their propagation throughout the brain [2, 4]. The glymphatic system is one such pathway; involved in clearance of metabolic waste products from the extracellular space in the central nervous system [10]. Predominantly active during sleep [11], this system is modelled on the inflow of cerebrospinal fluid (CSF) along perivascular spaces surrounding cerebral arteries, which then disperses throughout the brain interstitium via exchange with interstitial fluid (ISF), resulting in efflux of fluid and extracellular solutes via veinous perivascular space drainage out of the brain [10, 12]. Most compelling, lack of sleep is rapidly becoming recognised as a pathological driver of protein aggregation in neurodegenerative disease [13, 14], and sleep may act to enhance protein clearance from brain [12, 15], providing rationale for study of the function of this system in relation to clearance of ‘prion-like’ proteins from the brain [2].

Numerous independent lines of evidence, both post-mortem human [16, 17] and rodent studies [18–20], demonstrate the propensity of α-synuclein (αSyn) to propagate in a ‘prion-like’ mechanism throughout the brain, providing some explanation of the progressive and diffusive nature of neurodegenerative α-synucleinopathies such as Parkinson’s disease and dementia with Lewy bodies (DLB). As with other ‘prion-like’ proteins, this phenomenon is thought to be achieved by the initial release of an αSyn ‘seed’ into the extracellular space, transportation/transference into other cells, templating of intracellular fibrillar structures of αSyn and initiation of a self-amplifying cascade of accumulation, leading to further release of αSyn ‘seeds’ [21]. This characteristic of αSyn is now so well established, that there has been the recent suggestion by the pioneer of the prion field, Prof Stanley Prusiner, to abandon terms such as ‘prion-like’ in reference to neurodegenerative disease proteins such as αSyn, as the level of uncertainty as to their nature as prions is no longer justified [22]. Both cellular release and clearance mechanisms are thought to regulate the level of extracellular αSyn prions prone to propagation, hence αSyn clearance via the glymphatic system may represent a powerful determinant of propagation and progression of αSyn pathology in the brain.

Glymphatic perivascular fluid transport relies on the water channel aquaporin-4 (AQP4), which is predominantly expressed on the astrocytic endfeet that ensheathe cerebral vasculature [23, 24]. AQP4 deletion thus causes significantly suppressed CSF-ISF exchange and reduced clearance of β-amyloid [25, 26], tau [27–29] and αSyn [30, 31] from the brain. Not only is AQP4 expression essential, but its appropriate polarisation to astrocytic endfeet is required for efficient glymphatic and solute clearance [24, 28, 32–34]. Like the function of the glymphatic system, organisation of AQP4 in this way declines with age [35, 36], suggesting that impaired glymphatic function may contribute to the vulnerability of the aged brain to aberrant protein deposition, and as such, the onset of sporadic neurodegenerative disease. There is a growing body of evidence to suggest impaired glymphatic function in α-synucleinopathies such as Parkinson’s disease [37–43], but the causes of this are as yet unknown. For instance, it remains to be addressed how propagating αSyn pathology functionally affects the glymphatic system, and furthermore how altered glymphatic function affects αSyn dynamics and the ability of this prion protein to propagate across the central nervous system.

In this study, we make use of a mouse model of αSyn propagation, to better understand the effects of propagating αSyn aggregation on glymphatic function. We first measure glymphatic function and examine expression levels of AQP4 and its endfoot protein complex binding partners, both temporally and spatially in the brain. We then use a pharmacological approach to inhibit glymphatic function in this model setting, to determine whether impaired glymphatic function can accelerate or exacerbate αSyn aggregation and neuropathology in the brain.

## Materials and methods

### Generation of αSyn pre-formed fibrils

Recombinant human wildtype αSyn was expressed in BL21 (DE3) competent E.coli bacteria and purified with ion exchange and size-exclusion chromatography, dissolved in sterile phosphate buffered saline (PBS) and filtered sterilized (0.22 µm) before the concentration was adjusted to 5 mg/ml. Monomeric protein was converted to fibrils as per [44], by agitation at 1,000 RPM at 37 °C for 7 days in a ThermoMixer® C with ThermoTop® lid (Eppendorf). Pre-formed fibrils (PFFs) were aliquoted and stored at −80 °C until required. On the day of use, aliquots were thawed and diluted with sterile PBS to 2 mg/ml, and (where stated) sonicated to reduce the size of PFFs in solution as per [44]. Briefly, diluted PFF/monomer solutions were sonicated with a probe sonicator (Fisherbrand™ Model 50 Probe Dismembrator, Fisher Scientific) in 1 s on/off pulses for 60 s at 30% amplitude. Prepared monomer and PFF solutions were kept at 4 °C and room temperature respectively until needed.

### Validation of generated αSyn PFFs

#### Aggregate formation (sedimentation assay)

Formation of aggregates in the generated PFF solution was confirmed by sedimentation, similar to [45]. Briefly, PFF and monomer samples were diluted to 0.5 mg/ml in sterile PBS and centrifuged at 10,000 x *g* for 30 min at room temperature. Supernatants were collected and pellets were resuspended in the same volume of PBS. Samples were mixed 50:50 with 2x Laemlli buffer (Sigma), heated to 100 °C for 10 min, cooled on ice and then run in duplicate on a NuPAGE 4-12% Bis-Tris gel (ThermoFisher), for 30 min at 200 V in MES SDS running buffer in the presence of antioxidant (Bolt™ antioxidant, Invitrogen), using SeeBlue® Plus2 protein ladder (Invitrogen) (which contains a 14 kDa marker) to interpret molecular weight. Protein bands were stained by incubating the gel in Coomassie Brilliant Blue (Sigma) for 20 min at room temperature on an orbital shaker. The gel was then washed in distilled water (dH_2_O) before destaining overnight in destain solution (25% methanol, 5% acetic acid, 70% dH_2_O). The gel was washed again in dH_2_O prior to imaging on an ImageQuant™800 (Amersham) and bands analysed using ImageJ (v1.51j8).

#### Amyloid conformation (thioflavin-T fluorescence)

The presence of amyloid conformations in the generated PFF solution was confirmed by Thioflavin T (ThT) fluorescence, similar to [45]. Briefly, ThT reaction solutions were made by mixing ThT with either PFFs or monomers. 150 µl of solutions containing 12.5 µM ThT and 50 µg/ml of either PFFs or monomers in PBS were loaded in triplicate onto a black clear-bottomed 96-well plate. Well fluorescence was measured on an IVIS™ Spectrum imaging system with excitation at 430 nm and emission at 500 nm.

#### αSyn aggregate formation (dot blot)

Dot blots were used to confirm immunoreactivity of generated PFFs with a commonly used conformation-specific anti-αSyn antibody ([MJFR-14-6-4-2], Abcam). 1.5 µl of serially diluted monomer or (un)sonicated PFF samples were loaded onto nitrocellulose membrane (0.45 µm, Amersham) and left to dry overnight. Membrane was blocked in 5% non-fat milk in Tris-Buffered Saline with 0.1% Tween-20 (TBST) for 3 hrs at room temperature on an orbital shaker before being incubated in the primary antibody (1:1,000, conformation-specific anti-αSyn antibody [MJFR-14-6-4-2], Abcam) in 1% non-fat milk in TBST overnight at 4 °C. The membrane was then washed (3 x 5 min) in TBST and incubated in secondary antibody (1:5,000, goat anti-rabbit HRP-conjugated secondary, Abcam) in 1% non-fat milk in TBST for 3 hrs at room temperature. Membrane was washed again (3 x 5 min) in TBST before visualising immunoreactivity chemiluminescently (Pierce™ ECL Western Blotting Substrate, ThermoScientific), imaging on an ImageQuant™800 (Amersham) and quantifying signal intensity using ImageJ (v1.51j8).

#### Seeding capacity (kinetic ThT fluorescence)

The seeding capacity of PFFs was confirmed by their ability to decrease the time of monomer conversion to fibrils, monitored by ThT fluorescence, as per [44]. Reaction solutions were setup in PBS by mixing αSyn monomer starting material (0.5 mg/ml final concentration) with ThT (12.5 µM final concentration), and (un)solicited PFF (or monomer control) seeds (1:100 monomer to PFF ratio, 5 µg/ml final concentration). Solutions were loaded onto a black clear-bottomed 96-well plate in triplicate, three 1.5 mm glass beads added to each reaction well, and the plate sealed with optically clear plate-sealing film. Well fluorescence was measured immediately after setup using an IVIS™ Spectrum imaging system with excitation at 430 nm and emission at 500 nm, and then twice daily, for 4 days. When not being read, the plate was incubated in a ThermoMixer® C with ThermoTop® lid (Eppendorf) at 37 °C, and agitated at 600 RPM on a 1 min on/off schedule. Normalised fluorescence signal was fitted to sigmoidal curves, and Time_50_ values (time taken for monomers to aggregate and reach 50% maximal fluorescence with ThT) extracted for comparison of well conditions.

#### Propagation induction (*in vitro*)

The ability of PFFs to induce endogenous aggregation of αSyn *in vitro* was confirmed using SH-SY5Y cells overexpressing haemagglutinin (HA)-tagged human αSyn, similar to [46]. These cells were generated with a pcDNA3.1 plasmid and selected with G418 [47]. Proliferating overexpressing SH-SY5Y cells were cultured as previously described [48], passaged and resuspended in Neuralbasal media supplemented with B-27, glutaMAX and antibiotic/mycotic solution. Cells were seeded in this media supplemented with 30 μM retinoic acid and 10 ng/ml BDNF (R&D Systems) into polyornithine, fibronectin (2 μg/ml) and laminin (1 μg/ml) coated plates at 3 × 10^5^ cells/ml. Cells were differentiated for 4 days and then treated with either 2 μg/ml monomeric αSyn, unsonicated or sonicated PFFs for 3 days. Media was changed and cells incubated for a further 5 days, with a final media change 48 hrs before cell harvest. Cells were harvested with trypsin and washed in PBS. Cell pellets were lysed in 50 µl 1% (v/v) TX-100, 50 mM Tris, pH 7.5, 750 mM NaCl, 5 mM EDTA, 4 units RQ1 DNase (Promega), protease and phosphatase inhibitors (ThermoFisher) on ice for 20 min. Lysates were pelleted at 17,000⍰×⍰*g* for 20 min at 4 °C. TX-100 soluble fractions were placed in fresh tubes and protein concentration measured using the BCA protein assay. Insoluble pellets were solubilized in 50 µl 8 M urea, 2% (w/v) SDS, 10 mm Tris, pH 7.5, 4 units RQ1 DNase, protease and phosphatase inhibitors for 15 min at room temperature. Debris was removed by centrifugation at 17,000⍰×⍰*g* for 20 min.

Protein (5 µg) was loaded on NuPAGE 4–12% Bis-Tris gels (ThermoFisher), separated by electrophoresis and transferred to Hybond PVDF membrane (GE Healthcare). The amount of urea-SDS fraction loaded for each sample was calculated as a proportion of the protein concentration in the TX-100 soluble fraction. Because of its purity, β-actin was not detectable in the insoluble fraction, as would be expected. For αSyn blots, PVDF membranes were fixed with 4% paraformaldehyde (PFA) and 0.01% (v/v) glutaraldehyde for 30 min at room temperature. Membranes were blocked with 5% non-fat milk in PBS with 0.1% Tween-20, and incubated with the following primary antibodies: αSyn pS129 (abcam, ab51253), β-actin (abcam, ab6276), HA (Biolegend, HA.11). Following incubation with respective HRP-conjugated secondary antibodies, blots were incubated with Immobilon Luminata Forte enhanced chemiluminescence (Merck) and images acquired using Image Lab software (BioRad).

### Animals

Tg(Prnp-SNCA*A53T)83Vle were generated by Prof Virginia Lee [49] and were obtained from Jackson Laboratories (stock no.: 004479). Mice were hemizygously bred and homozygous offspring (henceforth referred to as M83 mice) were used for experiments at 2-5 months-of-age. Age-matched C75Bl6/J mice (henceforth referred to as wildtype mice) were used as controls and obtained Charles River, UK. Animals were housed on a 12-hr light/dark cycle in groups of two to five in individually ventilated cages with *ad libitum* access to food and water and provided with environmental enrichment (cardboard houses and tubes, paper nesting). Equal numbers of both male and female animals were used in experimental groups. All animal work was performed in accordance with the UK’s Animals (Scientific Procedures) Act of 1986 and approved by UCL’s internal Animal Welfare and Ethical Review Board.

### Intracerebral injection of αSyn

Mice (2 months-of-age) were anesthetized with 2% isoflurane in O_2_ at a delivery rate of 1 l/min and positioned in a stereotaxic frame in the horizontal skull position. Buprenorphine (Vetergesic, 0.1 mg/kg, subcutaneously) was administered for peri/post-operative analgesia, prior to making a midline incision on the top of the head to expose the skull. A burr hole was then made with a microdrill above the injection location (+⍰0.2 mm anteroposterior, and +⍰2 mm mediolateral to bregma). 2.5 µl/site of either sonicated αSyn PFFs or monomers (both at 2 mg/ml, 5 µg/site) were unilaterally infused at a rate of 0.5 µl/min into the striatum and overlying cortex (−1.8 and −⍰0.8 mm ventrodorsal to the brain surface respectively) using a 5 µl Hamilton syringe. After each infusion, the needle was left *in situ* for 5 min to prevent injectate reflux. The scalp was then sutured closed, and the animal left to recover in a heated recovery chamber until it had regained consciousness. It was then returned to its home cage, and its weight monitored daily for 7 days post-surgery to ensure complete recovery.

### mRNA and protein extraction and analysis

For time course expression studies, mice received intrastriatal/cortical injections with αSyn PFFs or monomers as above, and were left to age for the appropriate time period, after which animals were killed by overdose of sodium pentoparbital (10 ml/kg, intraperitoneally (i.p.)), the brain was removed from the skull and the ipsilateral and contralateral striatal and midbrain tissue was rapidly dissected out. Excised tissue was snap frozen immediately on dry ice and stored at −80 °C until further processing. For extraction, brain samples were homogenised by sonication and mRNA, protein and DNA were separated using TRIzol™ (Invitrogen)/chloroform centrifugation. mRNA was isolated from the aqueous phase using the PureLink™ RNA Mini Kit (Invitrogen) as per the manufacturer’s instructions, which yielded RNA with a 260/280 absorbance ratio of 2.03 (±0.04). RNA was converted to cDNA using the QuantiTect Reverse Transcription Kit (Qiagen) as per the manufacturer’s instructions, which was then quantified by qRT-PCR using TaqMan™ Universal PCR Master Mix and TaqMan™ probes against AQP4 (Mm00802131_m1), SNTA1 (Mm01251334_m1), DTNA (Mm00494555_m1), DMD (Mm00464475_m1) and DAG1 (Mm00802400_m1), normalising to GFAP (Mm01253033_m1) and using GAPDH (Mm99999915_g1) and ACTB (Mm02619580_g1) as housekeeper controls (all ThermoFisher), and the 2^-ΔΔCt^ method for analysis [50], normalising to wildtype baseline data.

Proteins were extracted from the phenol phase of the same brain samples following TRIzol™/chloroform centrifugation, as per the manufacturer’s instructions. Proteins (10 µg) were heated to 100 °C for 10 min with Laemlli buffer, cooled on ice and then run on a NuPAGE 4-12% Bis-Tris gel (ThermoFisher), for 30 min at 200 V in MES SDS running buffer in the presence of antioxidant (Bolt™ antioxidant, Invitrogen), using SeeBlue® Plus2 protein ladder (Invitrogen) to interpret molecular weight. Proteins were transferred onto nitrocellulose membrane (0.45 µm, Amersham), which was fixed with 4% PFA for 30 min before being blocked in 5% non-fat milk in TBST for 3 hrs (both incubations at room temperature). Membranes were then incubated in primary antibody (1:1,000 anti-αSyn pS129 (Abcam), or 1:1,000 anti-β-actin (Abcam)) in 1% non-fat milk in TBST overnight at 4 °C. Membranes were then washed (3 x 5 min) in TBST and incubated in secondary antibody (1:5,000, goat anti-mouse or rabbit anti-mouse HRP-conjugated secondaries, both Abcam) in 1% non-fat milk in TBST for 3 hrs at room temperature on an orbital shaker. Membranes were washed again (3 x 5 min) in TBST before visualising immunoreactivity chemiluminescently (Pierce™ ECL Western Blotting Substrate, ThermoScientific), imaging on an ImageQuant™800 (Amersham) and quantifying signal intensity using Image Lab software (BioRad).

### Cisterna magna infusion for assessment of glymphatic function

Glymphatic function was assessed in the mouse brain by infusion of a fluorescent tracer to the cisterna magna, followed by brain slice imaging for quantification of CSF-brain influx, similar to that previously published by our group [51]. Briefly, mice were anesthetized with 2% isoflurane in O_2_ at a delivery rate of 1 l/min and positioned in a stereotaxic frame with the head flexed to ∼50°. A midline incision was made at a midpoint between the skull base and the occipital margin to the first vertebrae. The underlying muscles were parted to expose the atlanto-occipital membrane and dura mater overlaying the cisterna magna, and a durotomy was performed using a 23-gauge needle. An intrathecal catheter (35–40 mm port length, 80–90 mm intravascular tippet length, Braintree Scientific, US) extended with polyethylene tubing (0.4 mm × 0.8 mm, Portex™, Smiths Medical) and attached to a 100 µl glass Hamilton syringe driven by a microinfusion pump (sp210iw syringe pump, World Precision Instruments, US) was filled with low molecular weight fluorescent tracer, Texas Red™-conjugated dextran (TxR-d3, 3 kDa; ThermoFisher) at 0.05% in filtered artificial CSF (Tocris), advanced 1 mm into the cisternal space, and sealed and anchored in place with cyanoacrylate. Once dry, TxR-d3 was infused via the implanted cannula (2 µl/min over 5 min). 30 min after the start of the infusion, mice were killed by overdose with sodium pentobarbital (10 ml/kg, i.p.) and the brain was quickly removed. Brains were then drop-fixed in 4% PFA for 48 hrs at 4 °C, prior to cryoprotection (30% sucrose in PBS for 72 hrs), freeze embedding in Optimal Cutting Temperature (OCT) compound, and sectioning (100 μm coronal sections) onto SuperFrost Plus™ microscope slides (VWR) using a cryostat (CM3050 S, Leica). Slides were imaged using an IVIS Spectrum imaging system (PerkinElmer) (Ex. 570 nm, Em. 600nm) and fluorescence signal quantified in the ipsilateral and contralateral striatum (∼+0.2 mm from bregma) and midbrain (∼-3 mm from bregma) using ImageJ (v1.51j8).

### Drug treatments

Glymphatic inhibition was achieved using TGN-020 (N-1,3,4-thiadiazol-2-yl-3-pyridinecarboxamide, Tocris Bioscience). Mice were treated i.p. with either TGN-020 (50 mg/kg in 20 ml/kg 0.9% sodium chloride (saline)) or vehicle (20 ml/kg saline). For acute studies, animals were given a single i.p. injection of either TGN-020 or vehicle. For chronic studies, animals were given i.p. injections of either TGN-020 or vehicle three times per week for 6 weeks (alternating the injection side each day), starting ∼3 days post-surgery.

### Assessment of acute αSyn clearance

Following acute treatment with TGN-020, αSyn PFFs were injected unilaterally into the striatum as described above (5 µg delivered in 2 mg/ml at 0.5 µl/min, +0.2 mm anteroposterior and +2 mm mediolateral to bregma and −1.8 mm ventrodorsal to the brain surface). The needle was left *in situ* while the cisterna magna was surgically exposed (also described above) and cleaned with an ethanol soaked swab. 15 min after the start of the striatal injection a durotomy was performed using a 23-gauge needle, allowing free-flowing CSF to be collected using a narrow bore pipette tip. Following collection, mice were killed by overdose with sodium pentobarbital (10 ml/kg, i.p.). The brain was removed from the skull and snap frozen immediately on dry ice and stored at −80 °C until further processing. The collected CSF was centrifuged briefly to pellet any red blood cell contamination. Spectrophotometric analysis (417 nm, NanoDrop ND-1000, Fisher Scientific) of hypotonically disrupted blood cell contaminants (freeze-thawing of rehydrated blood pellets) measured blood contamination to be <0.01% in all samples. Total tau concentration in CSF samples and homogenised (10% (w/v) in sterile PBS containing protease inhibitor cocktail, phosphatase inhibitor cocktails I and II (Sigma), at a final dilution of 1:100, and 1 mM phenylmethylsulphonyl fluoride) brain tissue was quantified by dot blot (described above) using a conformation-specific anti-αSyn antibody ([MJFR-14-6-4-2], Abcam), and diluted PFFs as a standard series. The total mass of αSyn left in the brain and the concentration of PFFs in extracted CSF were calculated.

### Open field task

The open field task was used to test for motor impairments. The apparatus consisted of a 45⍰×⍰45⍰×⍰45 cm white Perspex box open at the top. At the start of the test the mouse was placed in the centre of the field, and allowed to explore the apparatus for 5 min. The open field was thoroughly cleaned with 70% ethanol in between trials. Each mouse was recorded using a digital video camera (Konig, CASC300) placed approximately 50 cm above the apparatus. Animals’ paths were tracked using AnimalTracker (http://animaltracker.elte.hu/), which automatically determined the distance travelled and time spent in the central and peripheral areas of the field. Path tracking analysis was carried out with the experimenter blind to experimental/treatment group.

### Magnetic resonance imaging

At the end of drug treatment studies (6 weeks post-αSyn injection), structural 3D MR images of mouse brains were acquired. For this, mice were anesthetized with 2% isoflurane in a mixture of air (0.4 l/min) and O_2_ (0.1 l/min), and positioned into an MR-compatible stereotaxic frame, with 1.5% isoflurane delivered (in the same gas mixture) via a nose cone. A horizontal-bore 9.4T Bruker preclinical system (BioSpec 94/20 USR, Bruker) was used for acquisition of isotropic 3D structural T_2_-weighted MR images of the entirety of the mouse brain, using a volume coil and 4-channel surface coil for radiofrequency transmission and signal detection, respectively. A fast spin echo sequence with the following parameters was used: FOV⍰=⍰16⍰×⍰16⍰×⍰16 mm, matrix⍰=⍰128⍰×⍰128⍰×⍰128, repetition time⍰=⍰700 ms, echo time⍰=⍰40 ms, 2 averages.

For analysis, ITK-SNAP (v4.2.0) [52] was used for manual segmentation of the striatum and midbrain, referring to the Allen Mouse Brain Reference Atlas (https://mouse.brain-map.org/static/atlas) for anatomical guidance. Segmentation was primarily performed in the coronal plane, with adjustment of regions-of-interest (ROIs) boundaries in the other two imaging planes. ROI pixel numbers were converted to volumes which are presented by multiplying by the pixel volume (0.001953125 mm^3^).

### Immunohistochemistry

At the end of chronic treatment experiments, CSF samples were extracted from the cisterna magna for quantification of αSyn content (as described above) prior to perfuse fixation of mice. For this, mice were injected with an overdose of sodium pentobarbital (10 ml/kg, i.p.) and transcardially perfused with approximately 10 ml PBS immediately followed by the same volume of 4% PFA. Brains were extracted from the skull and drop-fixed in 4% PFA for 24 hrs at 4 °C. Brains were then cryopreserved (30% sucrose in PBS for 72 h at 4 °C), freeze embedded in OCT, and serial coronal sections collected (50 μm thick) onto SuperFrost Plus™ microscope slides (VWR) using a cryostat (CM3050 S, Leica). Slides were stored at −20 °C until further processing. On the day of staining, sections were brought to room temperature before being baked at 60 °C for 60 min. Antigen retrieval was performed by incubation of tissue slides in high pH eBioscience™ antigen retrieval solution (Invitrogen) with mechanical compression under teflon [53] for 30 min in a steamer. Slides were then blocked in 10% normal goat serum in PBS with Triton-X (0.2%) (PBST) for 2 hrs at room temperature before incubation with primary antibody (1:500, conformation-specific anti-αSyn antibody [MJFR-14-6-4-2], Abcam) in blocking solution overnight at room temperature. Sections were washed (3 x 5 min) in PBST before incubation in anti-rabbit secondary antibody conjugated to Alexa Fluor™ 568 (1:800, donkey, Invitrogen) for 2 hrs in the dark at room temperature. Sections were then washed again (3 x 5 min) in PBST and mounted under coverslips using Fluoromount-G™ with DAPI (Invitrogen). Slides were kept in the dark until imaging.

Single plane, tiled images were captured on a fluorescence microscope (Leica DMi8) using a 10⍰×⍰1.3 NA objective. For analysis, using ImageJ (v1.51j8), a ROI was drawn on images encompassing the striatum in at least 6 sections from each brain, with the experimenter blind to animal experimental/treatment group. Measures of % area coverage (after thresholding for immunoreactivity) were extracted from each and averaged across slides analysed from each brain.

### Statistical analysis

All data is presented as group means ± standard error of the mean (SEM), with individual datapoints shown where possible. For comparisons between groups, either 1-way or 2-way Analysis of Variance (ANOVA) was used, depending on the dataset (ANOVA type denoted on individual graphs), with either Tukey’s multiple comparisons tests for individual comparisons, or Šidák’s or Bonferroni multiple comparisons tests for grouped data comparisons. For XY plots, simple linear regression analyses were used to test for correlations. For the kinetic ThT assay, data was fitted to a sigmoidal four parameter curve, extracting 50% maximal (Time_50_) values. All graphs were constructed, and statistical comparisons made using GraphPad Prism (v10.3.0).

## Results

### αSyn PFFs exhibit amyloid characteristics and induce propagation *in vitro* and *in vivo*

Firstly, amyloid αSyn PFFs were generated, from recombinant human wildtype αSyn by agitation [44, 54], for subsequent injection and induction of pathological αSyn propagation *in vivo*. In order to confirm the amyloid characteristics and ability of the generated PFFs to induce αSyn propagation prior to injection, a number of quality control experiments were conducted.

Successful conversion of monomers to fibrils was firstly confirmed by sedimentation. Following centrifugation, significantly greater protein was observed in the pellet compared to supernatant fractions of PFF samples (p<0.0001). The inverse was observed in the monomeric starting material (p<0.0001) (**Fig. 1A, B**), indicating successful conversion to fibrils. The presence of amyloid structures in generated PFFs was then confirmed by ThT fluorescence. Significantly greater fluorescence was observed when ThT was incubated with PFFs compared to monomers and all other controls (p<0.01 in all comparisons) (**Fig. 1C**). In order to confirm the presence of αSyn aggregates in generated PFFs, dot blot analysis was conducted using an antibody against conformation-specific αSyn ([MJFR-14-6-4-2], Abcam) (**Fig. 1D**). Increasing immunoreactivity with the amount of protein loaded was observed in both sonicated and unsonicated PFFs samples, but not with monomers (**Fig. 1D, E**). The ability of the generated PFFs to seed aggregation was then assessed, by agitation of αSyn monomers with PFF samples (1:100 PFF to monomer ratio) in the presence of ThT. Unsonicated PFFs reduced the time taken for monomers to aggregate and reach 50% maximal fluorescence with ThT (Time_50_) by 6.77 hrs compared to monomers alone (p<0.0001, inset, **Fig. 1F**). Sonication of PFFs prior to seeding further reduced Time_50_; 11.66 hrs quicker than monomers alone (p<0.0001, inset, **Fig. 1F**), indicating the significant seeding capacity of the sonicated PFFs generated.

**Figure 1.**
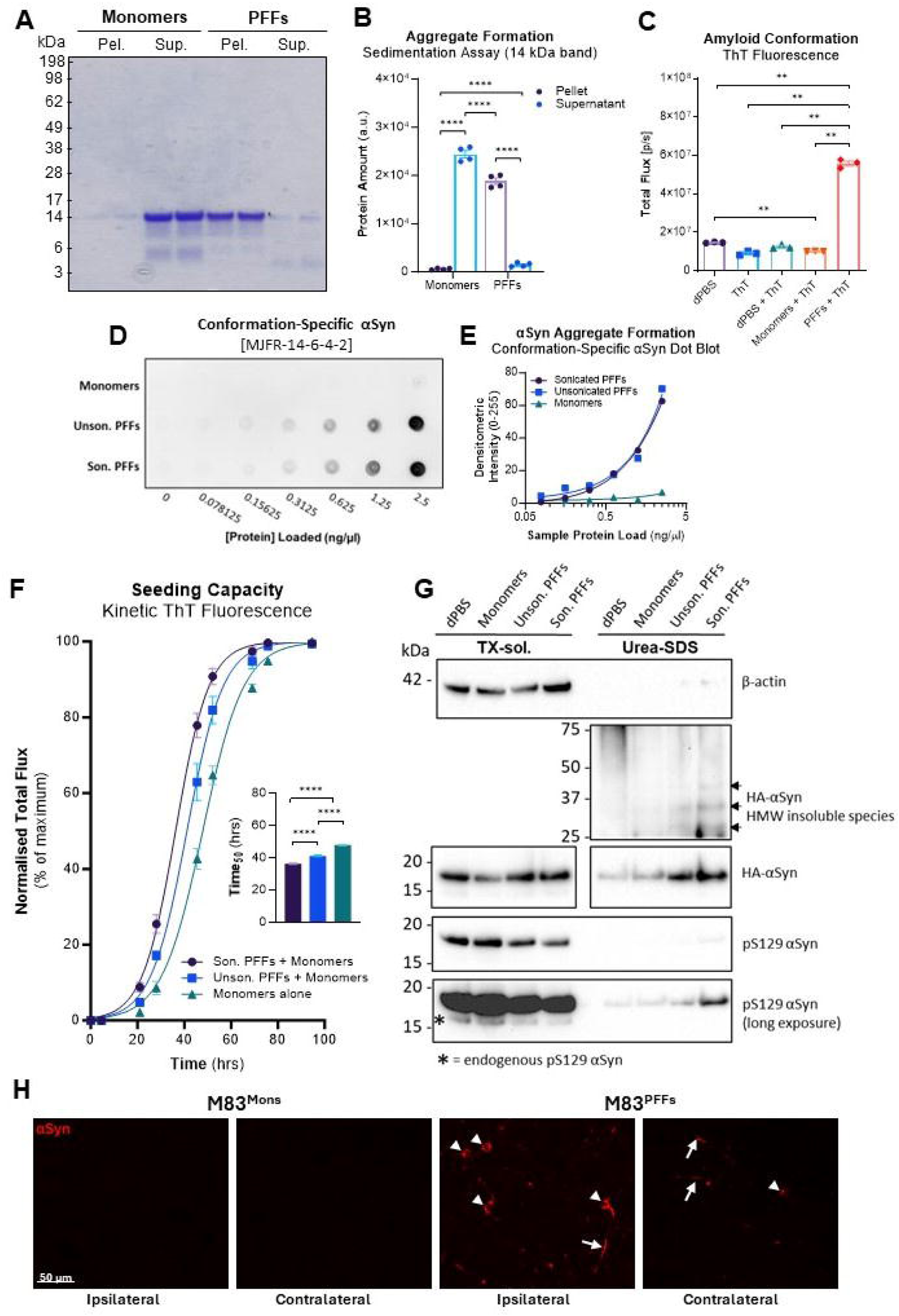
αSyn PFFs exhibit amyloid characteristics and induce propagation *in vitro* and *in vivo*. Aggregates are present in generated PFFs, as is evident by their sedimentation following centrifugation. (**A**) Coomassie stained electrophoresis gel showing that the majority of the protein (14 kDa, αSyn) is in the supernatant of the monomer starting material, but in the pellet of generated PFFs following centrifugation. Results from this assay are shown graphically in (**B**). Generated PFFs fluoresce with ThT (**C**) indicating the presence of amyloid structures. Both sonicated and unsonicated PFFs are also immunopositive for a commonly used conformation-specific αSyn antibody (MJFR-14-6-4-2) (**D**, **E**), indicating the presence of aggregated αSyn protein in generated PFFs. Sonicated and unsonicated PFF also induce a stepwise left-ward shift in the sigmoidal intensity profile of ThT monitored αSyn aggregation *in vitro* (**F**), resulting in reduced Time_50_ values in PFF seeded samples compared to aggregate formation of monomers alone (**F, inset**). PFFs were also shown to induce propagation of αSyn *in vitro* in a cell culture system. Incubation of SH-SY5Y cells over expressing HA-tagged αSyn caused the increased accumulation of insoluble pS129 αSyn, aggregation of endogenously expressed (HA-tagged) αSyn and the occurrence of higher molecular weight (HMW) αSyn species (**F**, arrows). (**H**) Immunofluorescence images of αSyn pathology (aggregate conformation-specific) in the striatum (both ipsilateral and contralateral) of monomer and PFF injected M83 mice, illustrating both intracellular αSyn accumulations (arrowheads) and αSyn neurite pathology (arrows) in PFF injected but not monomer injected animals. Results from statistical tests (C and F inset, 1-way ANOVA; B, 2-way ANOVA) indicated with asterisks: **=p<0.01, ****=p<0.0001.

In order to confirm the seeding capacity of PFFs *in vitro* ahead of injection, SH-SY5Y cells overexpressing αSyn with a hemagglutinin (HA) tag [47] were differentiated for 4 days and then treated with either 2 μg/ml monomeric αSyn, unsonicated or sonicated PFFs for 3 days. Media was changed and cells incubated for a further 5 days, with a final media change 48 hrs before neurons were harvested. The majority of αSyn remained in the TX-100 soluble fraction of cell lysates, however treatment with sonicated PFFs increased the level of TX-100 insoluble (solubilized with urea-SDS) αSyn, which was phosphorylated at Ser129 (**Fig. 1G**). Note that in the insoluble fraction, β-actin was barely detectable, indicating the purity of the fraction, which is important as contamination with the soluble fraction and its much greater αSyn protein levels would limit the interpretation of results. Importantly, treatment with sonicated PFFs increased the level of HA-αSyn, indicating propagated accumulation of endogenously expressed αSyn in cells (**Fig. 1G**). There was also evidence of higher molecular weight (HMW) αSyn species in the insoluble lysate fraction following treatment with sonicated PFFs (arrows, **Fig. 1G**), indicating PFF induced αSyn aggregation.

Lastly to confirm seeding capacity and the ability of generated PFFs to induce propagation of αSyn pathology i*n vivo*, homozygous M83 mice (2 months-of-age) were injected with either monomers or sonicated PFFs into the left striatum and overlying cortex, as previously described [54], and left to age for 6 weeks. Immunofluorescence staining for αSyn aggregation ([MJFR-14-6-4-2], Abcam), revealed substantial intracellular αSyn accumulations (**Fig. 1H**, arrowheads) and αSyn-positive neurite processes (**Fig. 1H**, arrows) in both the ipsilateral, and also the contralateral striatum of PFF injected mice, but not in monomer injected animals. Together, these data indicate successful generation of αSyn PFFs from recombinant human wildtype protein, capable of pathological propagation induction *in vivo*.

### αSyn PFF seeding causes dystrophin associated complex dysregulation *in vivo*

For study of glymphatic mechanisms, cohorts of homozygous M83 mice (2 months-of-age) were injected with either monomers or sonicated PFFs into the left striatum and overlying cortex to initiate αSyn propagation *in vivo*, as above and as previously described [54]. Seeding with PFFs in this way resulted in significant decline of striatal AQP4 and expression of elements of the AQP4 dystrophin-associated complex (DAC) (**Fig. 2C**) compared to monomer injected controls, as early as 2 weeks post-injection (58.7% expression reduction on average in PFF injected mice compared to wildtype baseline, vs. 14.0% expression increase on average in monomer injected mice at the same timepoint, p<0.0001, **Fig. 2A, B**). This decline in AQP4 and DAC element expression in PFF injected mice continued to develop with time (66.6% and 76.3% expression reductions on average at weeks 4 and 6 post-injection, respectively, p<0.0001 compared to monomer injected controls at both timepoints, **Fig. 2A, B**). Most interestingly, when DAC mRNA expression profiles were correlated to phosphorylated (at Ser129) αSyn protein load, negative associations were observed; reduced DAC element expression with increased αSyn load. The strength of these correlations reached significance in elements of the DAC most proximal to the water channel (**Fig. 2C**) (AQP4, p=0.0294, and SNTA1, p=0.01, **Fig. 2D**) and approached significance in the distal elements (DTNA, p=0.0533, DMD, p=0.0805, and DAG1, p=0.0598, **Fig. 2D**), suggesting that αSyn deposition and AQP4 complex integrity are in some way linked.

**Figure 2.**
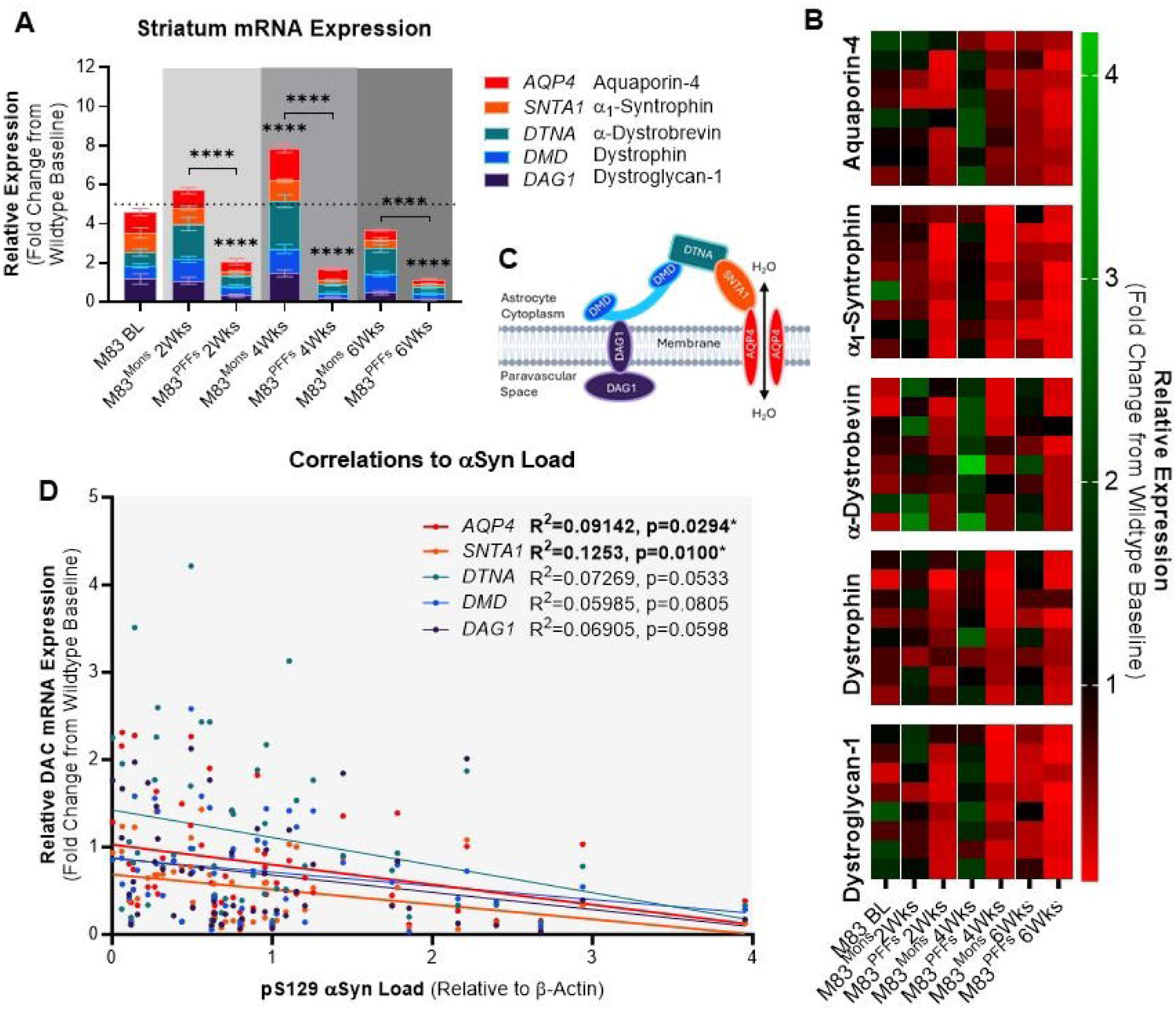
αSyn PFF seeding causes dystrophin associated complex dysregulation *in vivo*. (**A**) Striatal mRNA expression levels of AQP4 and DAC elements in experimental groups injected with either αSyn monomers or PFFs and left to age for 2, 4 or 6 weeks. Expression data is normalised to wildtype baseline expression levels (dotted line). Individual animal datapoints (n=8) shown in heatmap (**B**). (**C**) diagrammatic illustration of the DAC at the astrocytic endfoot membrane, with elements colour-coded to reflect data in (A). Correlation analysis of mRNA expression profiles with animal-by-animal αSyn load data, illustrating significant negative correlations with AQP4 and SNTA1, and correlations approaching significance in the other three DAC proteins. Results from statistical tests (B, 2-way ANOVA; D, simple linear regression fitting) indicated with asterisks: *=p<0.05, ****=p<0.0001. Asterisks in (A) indicate significance from M83 baseline (BL), asterisks with bars indicate timepoint comparisons.

This was not the case in the midbrain however; no noticeable correlations were observed between DAC mRNA expression and αSyn load (all linear regression R_2_ values <0.04, all p values >0.15, **Supp. Fig. 1D**). This perhaps is not surprising, given that this rostral region of the brain receives αSyn only towards the later stages of pathological development in this model [54]. Consistently, at week 6 post-injection, reduced AQP4 complex expression was observed in the midbrain of PFF injected mice (29.6% expression reduction on average in PFF injected mice compared to wildtype baseline levels, p<0.0001 compared to transgenic baseline expression, **Suppl. Fig. 1A, B**), suggesting that propagated αSyn deposition may directly affect AQP4 complex integrity, despite no overall correlation existing in this brain region.

### αSyn arrival coincides with glymphatic upregulation in the midbrain of PFF seeded mice

To investigate this suggestion further, we then assessed glymphatic function in this animal model. To our surprise, at weeks 4 and 6 post-injection, we observed enhanced bilateral tracer uptake in the midbrain of PFF injected mice following cisterna magna infusion, indicative of increased CSF-ISF exchange (p=0.0020 and p=0.0062 compared to transgenic baseline at weeks 4 and 6 respectively, **Fig. 3A, B**). This upregulation in PFF injected mice was significantly increased compared to monomer injected controls only at week 4 (p=0.0403), but not at week 6 (p=0.4615) however (**Fig. 3A, B**). Quantification of αSyn deposition in this same brain region revealed that this glymphatic upregulation coincides with a surge in αSyn accumulation (p=0.0353 compared to respective timepoint levels in monomer injected controls, **Fig. 3C, D**), suggesting that perhaps glymphatic function is upregulated in response to the arrival of propagated αSyn at this timepoint. In contrast, no corresponding change in tracer uptake was observed in the striatum of PFF injected animals, despite direct injection of αSyn in this region (**Suppl. Fig. 2**). Glymphatic upregulation compared to transgenic baseline in the midbrain however persists to week 6 (p<0.01 compared to transgenic baseline levels, **Fig. 3A, B**), resulting in normalisation of deposited αSyn (**Fig. 3C, D**), despite downregulation of AQP4 and DAC element expression in this region at this timepoint (**Supp. Fig. 1A, B**).

**Figure 3.**
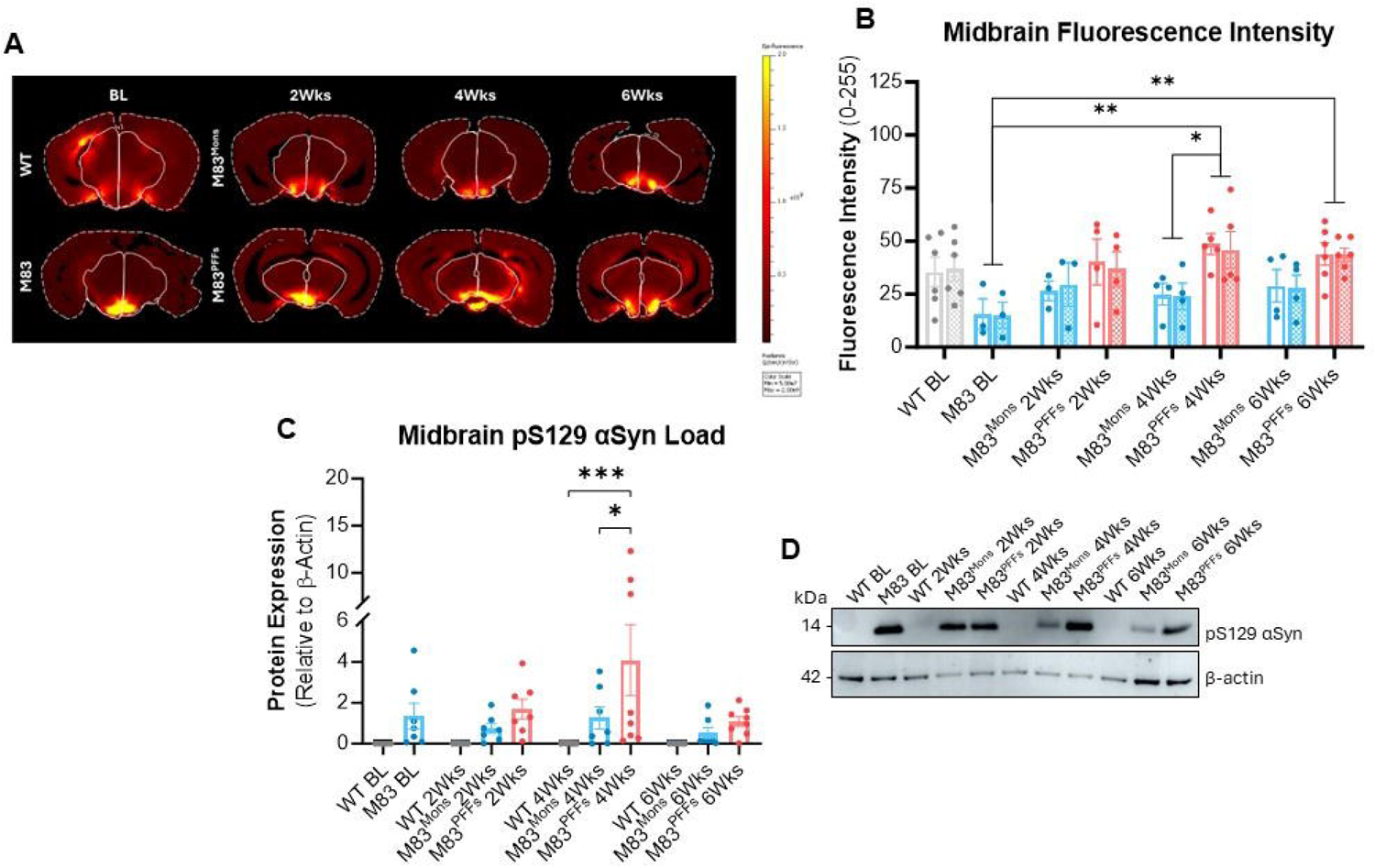
αSyn arrival coincides with glymphatic upregulation in the midbrain of PFF seeded mice. (**A**) Fluorescence intensity in the midbrain (solid outline) in brain sections (dashed outline) following cisterna magna infusion of Texas Red™-conjugated dextran (3 kDa). Intensity is quantified and graphed in (**B**) illustrating increased midbrain uptake at 4 and 6 weeks post-PFF injection. The ipsilateral midbrain area is illustrated with empty bars, the contralateral midbrain area is illustrated with hashed bars. (**C**) Quantification of αSyn at these same timepoints showed a surge of αSyn deposition at week 4, which resolves by week 6 after PFF injection. (**D**) Representative blots for data presented in (C). Results from statistical tests (B, 2-way ANOVA; C, 1-way ANOVA) indicated with asterisks: *=p<0.05, **=p<0.01, ***=p<0.001.

### Acute TGN-020 treatment reduces brain to CSF αSyn clearance

To investigate the relationship between glymphatic function and αSyn further, a pharmacological approach was taken for glymphatic inhibition; experimentally altering glymphatic function to determine the resultant effects on αSyn clearance from the brain. We have previously shown that acute TGN-020 treatment causes marked inhibition of glymphatic function [27]. Here, mice were systemically treated with either TGN-020 or vehicle prior to intrastriatal injection of either PFFs or aCSF. 15 min later, brain and CSF (from the cisterna magna) were collected for quantification of αSyn content by dot blot (**Fig. 4A, B**). TGN-020 treatment resulted in significantly greater αSyn retention in the brain compared to vehicle treatment (p=0.0492, **Fig. 4C**). Correspondingly, in the same mice, significantly reduced CSF αSyn was seen in drug treated mice compared to that observed in vehicle treated animals (p=0.0358, **Fig. 4D**). Together, these changes are consistent with glymphatic mediated clearance of αSyn.

**Figure 4.**
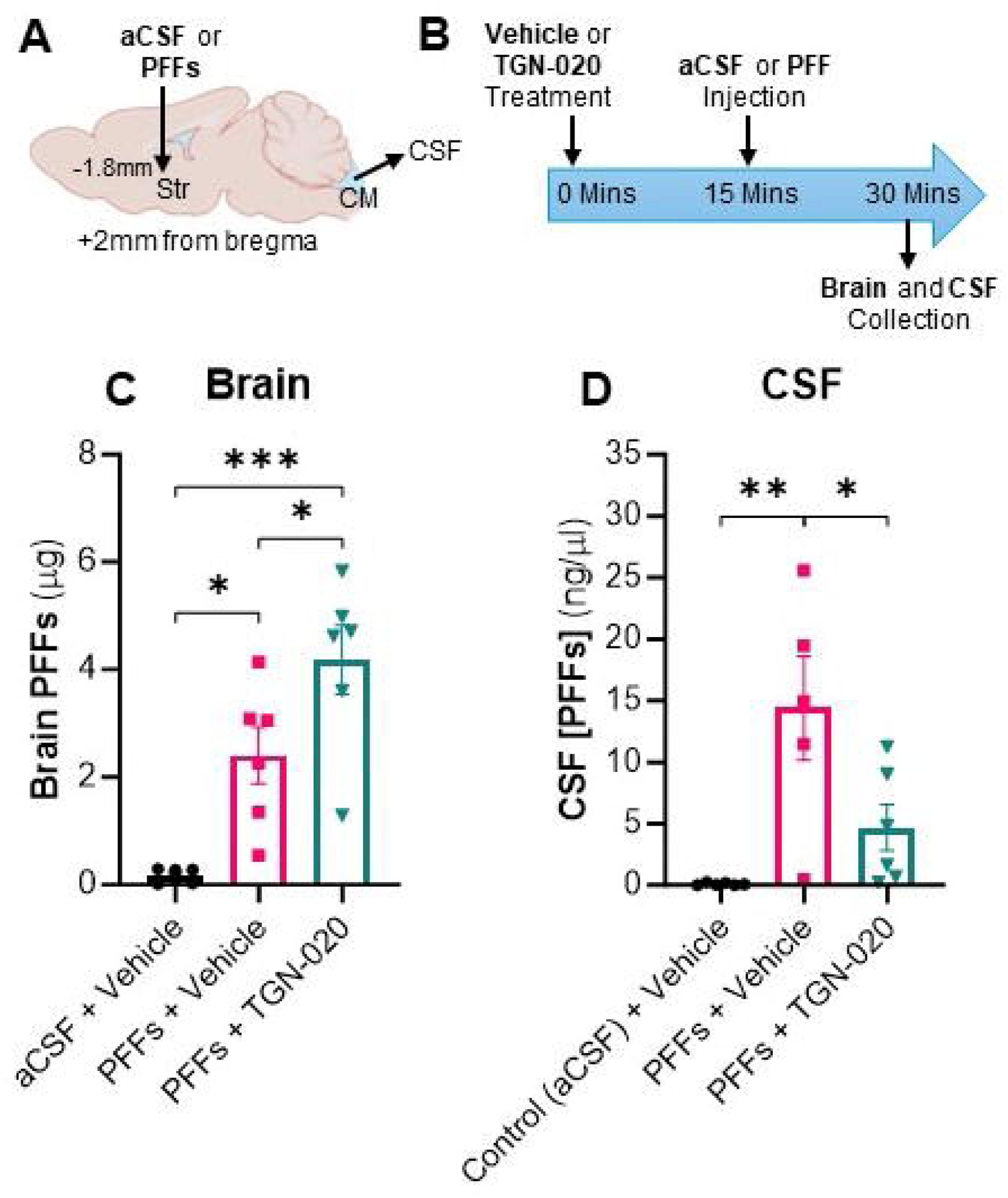
Acute TGN-020 treatment reduces brain to CSF αSyn clearance. (**A**) Diagrammatic representation and (**B**) time course of acute experiments, illustrating intrastriatal injection of aCSF or PFFs and pre-treatment with vehicle or TGN-020 (50 mg/kg). TGN-020 results in elevated levels of PFF brain retention (**C**) and a corresponding reduction in clearance to CSF (**D**). Results from statistical tests (1-way ANOVAs) indicated with asterisks: *=p<0.05, **=p<0.01, ***=p<0.001.

### Chronic TGN-020 treatment exacerbates pathology in the PFF-M83 model of αSyn propagation

We have previously shown that chronic treatment with TGN-020 results in sustained glymphatic inhibition [51], and saw above that acute glymphatic inhibition resulted in reduced clearance of αSyn from the brain. Therefore in order to determine whether a change in glymphatic function could affect disease progression *in vivo* in a disease model setting, as above, M83 mice (2 months-of-age) were injected with either monomers or PFFs (**Fig. 5A**) to initiate αSyn propagation [54], and subsequently treated chronically with either TGN-020 or vehicle (**Fig. 5B**). Motor behaviour of mice was assessed by open field at baseline (prior to intracerebral injection) and after 6 weeks of treatment, and structural MRI scans were acquired at the end of the experiments prior to brain and CSF collection for downstream analysis (**Fig. 5B**).

**Figure 5.**
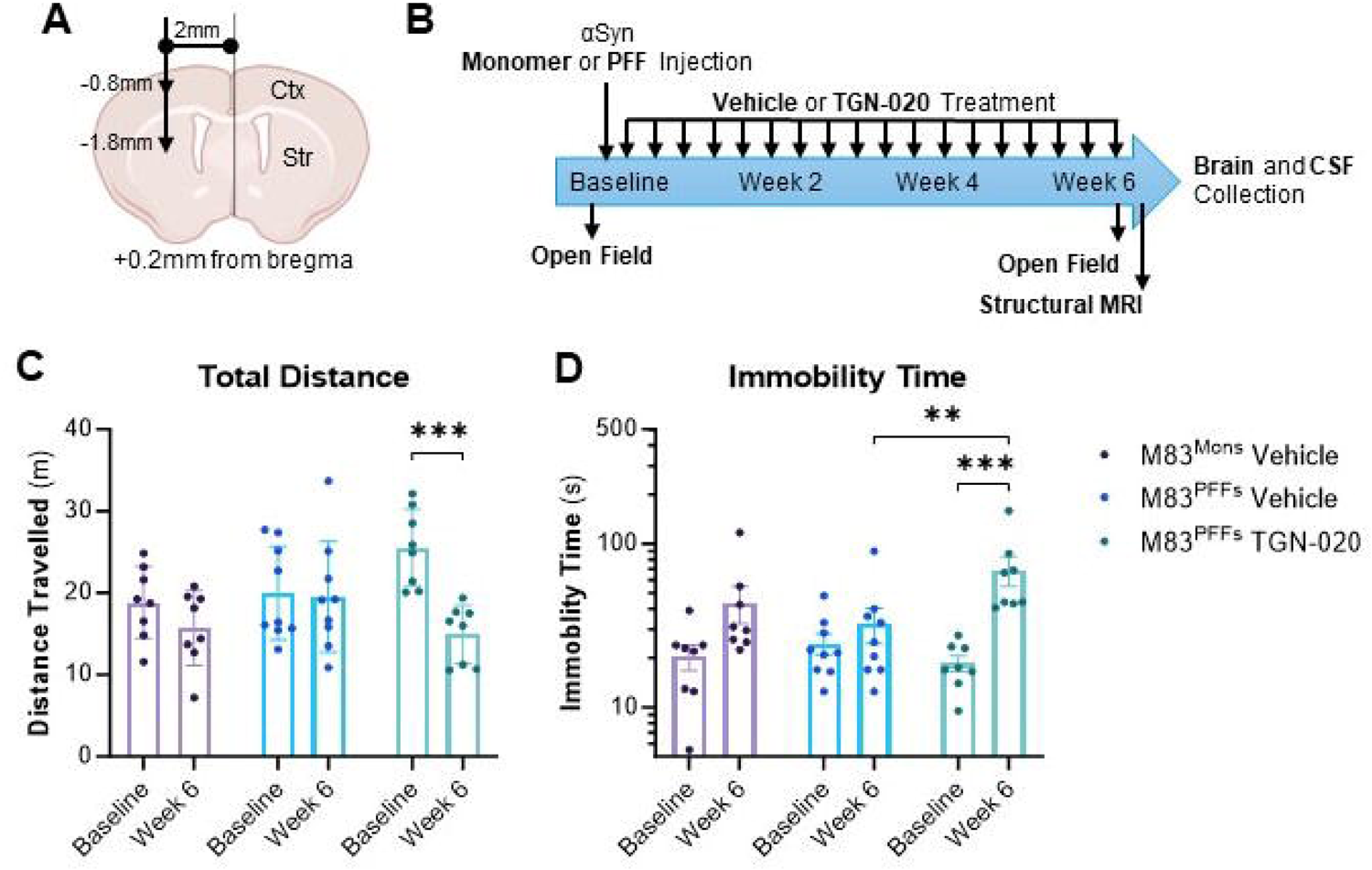
Chronic TGN-020 treatment induces behavioural deficits in the PFF-M83 model of αSyn propagation. (**A**) Diagrammatic representation and (**B**) time course of chronic experiments, illustrating location of intracerebral injection (striatum and overlying cortex) of either monomers or PFFs, followed by chronic treatment with TGN-020 (50 mg/kg, 3 times/week) for 6 weeks, with behavioural test and structural MRI acquisition timepoints noted. TGN-020 treatment resulted in reduced distance travelled in an open field arena after 6 weeks of treatment compared to baseline assessment (**C**), and increased immobility (**D**). Results from statistical tests (2-way ANOVAs) indicated with asterisks: **=p<0.01, ***=p<0.001.

TGN-020 treatment induced a locomotor phenotype in PFF seeded M83 mice. Drug treated animals exhibited significantly reduced distance travelled in the open field arena at week 6 compared to their baseline assessment (p=0.0005, **Fig. 5C**). This reduced locomotion of TGN-020 treated mice resulted in a significant increase in the time spent immobile in the arena, compared to both their own baseline assessment (p=0.0003, **Fig. 5D**), and also the vehicle treated PFF seeded group at week 6 (p=0.0069, **Fig. 5D**). No such differences were observed in monomer injected animals, or in PFF injected vehicle treated mice (**Fig. 5C, D**). In order to confirm that the observed impaired motor function was specific to transgenic mice seeded with PFFs and not a more general affect of drug treatment, experiments were repeated in wildtype PFF injected animals. No motor deficits were observed in either TGN-020 or vehicle treated PFF seeded wildtype animals (**Suppl. Fig. 3**), suggesting that the effect was unique to PFF seeded M83 mice, in which substantial αSyn propagation and deposition is observed (**Fig. 1H**).

Structural, anatomical MRI scans of mouse brains were acquired following 6 weeks of drug/vehicle treatment. In both PFF injected groups, significant atrophy was observed in the ipsilateral striatum compared to contralateral hemispheric volumes (p=0.0012 and p=0.0001, respectively, **Fig. 6A**). This interhemispheric difference was greater in TGN-020 treated mice however (16.27% in TGN-020 treated, and 12.80% in vehicle treated groups), and significantly reduced striatal volumes were observed in TGN-020 treated PFF injected mice compared to monomer injected vehicle treated controls (**Fig. 6A**). In the midbrain, significantly reduced volumes were observed in TGN-020 treated PFF injected mice compared to both PFF injected (p=0.0072) and monomer injected (p<0.0001) control mice (**Fig. 6B**). And TGN-020 treated animals were the only group in which significant ipsilateral atrophy was observed compared to contralateral volumes (p=0.0164, **Fig. 6B**).

**Figure 6.**
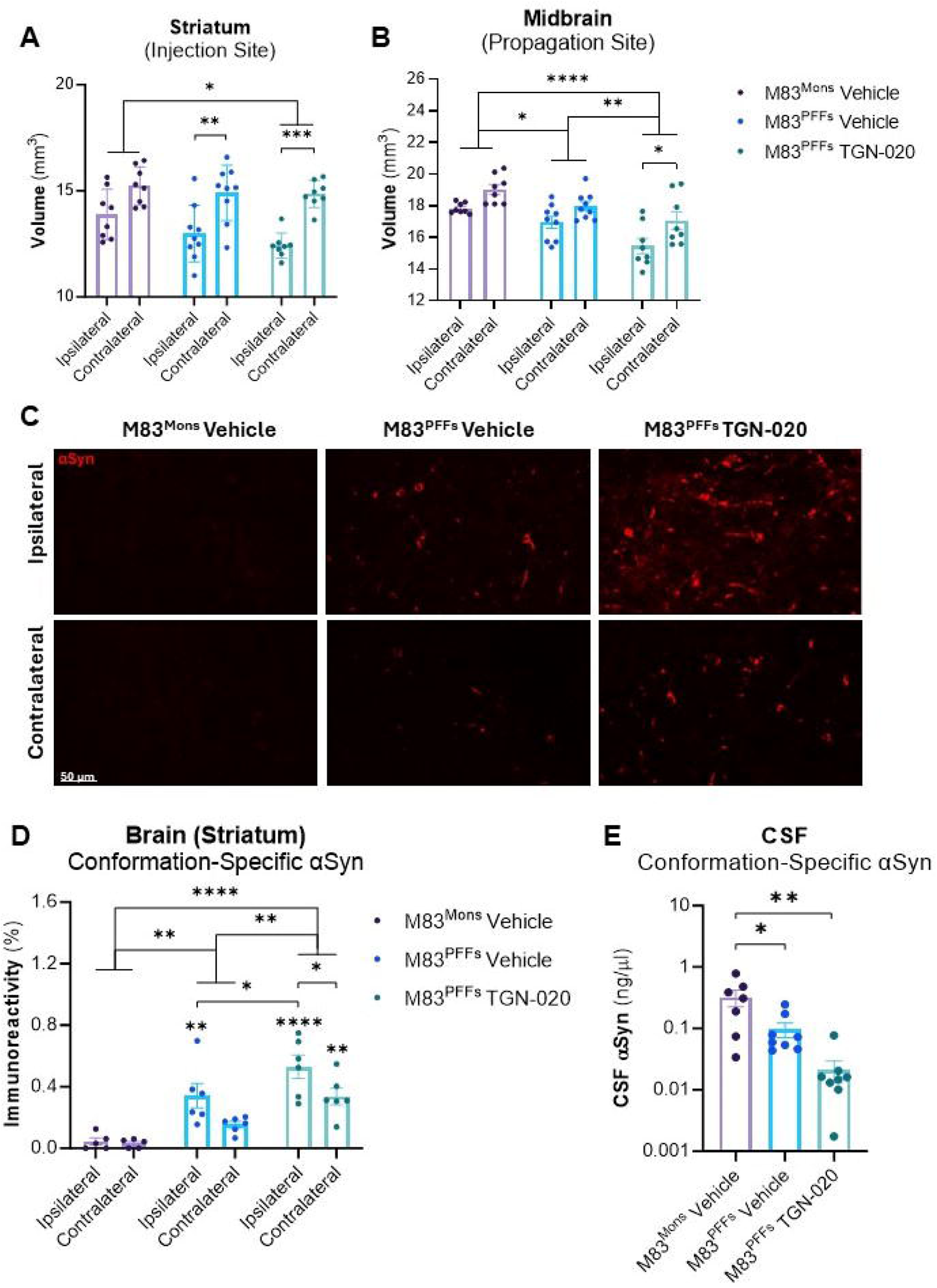
Chronic TGN-020 treatment exacerbates pathology in the PFF-M83 model of αSyn propagation. Volumes of the (**A**) striatum and (**B**) midbrain in chronically treated mice, indicating exacerbation of αSyn PFF induced atrophy following chronic treatment with TGN-020 for 6 weeks. (**C**) Representative immunofluorescence images of αSyn pathology (aggregate conformation-specific) in the striatum of treated mice, quantified and presented in (**D**), illustrating increased αSyn pathology with TGN-020 treatment. A corresponding decrease of CSF αSyn was observed in experimental animals. Results from statistical tests (A, B and C, 2-way ANOVAs; D, 1-way ANOVA) indicated with asterisks: *=p<0.05, **=p<0.01, ***=p<0.001, ****=p<0.0001. Asterisks without bars indicate individual hemispheric differences compared to the M83^Mons^ Vehicle group.

Following MRI acquisition, CSF samples were taken and brains collected for histology. Significantly greater levels of aggregated αSyn were observed in the striatum of TGN-020 treated PFF injected mice compared to both vehicle treated PFF injected (p=0.0048) and monomer injected (p<0.0001) controls (**Fig. 6C, D**). Furthermore, in the ipsilateral striatum, significantly greater αSyn was seen in TGN-020 treated mice compared to both PFF injected (p=0.0499) and monomer injected (p<0.0001) vehicle treated animals (**Fig. 6C, D**). Exacerbated αSyn deposition in the ipsilateral striatum of drug treated mice also resulted in a significant interhemispheric difference in these animals (p=0.0492, **Fig. 6C, D**). Similar to what was observed following acute TGN-020 treatment (**Fig. 4**), exacerbated αSyn load in the brain as a result of chronic drug treatment was accompanied by decreased CSF αSyn content: 90.33% reduction in PFF injected vehicle treated mice compared to monomer injected controls, and a further reduction of 7.53% (97.87% overall) in TGN-020 treated PFF injected mice compared to the monomer injected group (**Fig. 6E**).

## Discussion

In the current study we used a mouse model of αSyn propagation to investigate how impaired glymphatic function influences αSyn propagation dynamics in the brain and the effect of propagating αSyn itself on glymphatic function. We show that inducing αSyn pathology *in vivo*, by injection of αSyn PFFs into the striatum, leads to a robust reduced expression profile of the AQP4 endfoot protein complex, which negatively correlates to phosphorylated αSyn load. Conversely, propagation of αSyn to the midbrain with time inspires an enhancement of glymphatic function, which appears to manage the influx of propagating αSyn and normalises protein burden. To directly examine the influence of the glymphatic system on αSyn propagation and deposition, we then used a pharmacological approach to inhibit glymphatic clearance. Acute glymphatic inhibition reduced clearance of αSyn from brain to CSF, and extended inhibition of glymphatic function in PFF seeded mice led to exacerbated αSyn pathology, neurodegeneration, and motor behavioural deficits. Together, our data show that αSyn clearance and propagation are regulated by glymphatic function and suggest that AQP4 complex dysregulation may contribute to glymphatic impairment associated with Parkinson’s disease.

Recent years has seen a surge in research publications suggesting impaired glymphatic function in α-synucleinopathies such as Parkinson’s disease [37–43]. This body of work largely uses diffusion tensor imaging (DTI) and the ‘DTI along the perivascular space’ (DTI-ALPS) index as a non-invasive marker of glymphatic function, by isolating and measuring water diffusivity within perivascular spaces running parallel to deep medullary veins. The validity of DTI-ALPS as a marker for glymphatic function has recently been called into question [55], but given the wealth of studies now highlighting the potential involvement of the glymphatic system in this class of neurodegenerative disease, further work is justified to ascertain the precise mechanisms underlying the contribution of glymphatic dysfunction to disease progression in α-synucleinopathy.

The relationship between neurodegenerative disease pathology and perivascular AQP4 has been explored in Alzheimer’s [56], but the involvement of glymphatic clearance in α-synucleinopathies remains largely under researched. Hoshi and colleagues examined AQP4 expression in the temporal neocortex of patients with Parkinson’s disease and found a negative correlation between αSyn pathology and AQP4 expression [57]. We saw a similar relationship in the current study, in the striatum of mice seeded with αSyn, extending the observation to DAC proteins. To our knowledge however, AQP4 polarisation is yet to be quantified in the Parkinson’s disease brain. But consistent with what has been shown post-mortem, our data suggests that not only is AQP4 affected in pathology, but so too is the complex of proteins which anchors the water channel at the membrane, facilitating the glymphatic clearance of solutes, including αSyn, from the brain.

There has been more work looking at the relationship between glymphatic function and αSyn pathology in the mouse brain however. Cui and colleagues showed that when AQP4 expression was reduced (heterozygous AQP4 knockout mice), αSyn pathology, dopaminergic neurodegeneration, and motor behavioural impairments induced by injection of αSyn PFFs were exacerbated [30]. This study was recently added to by Zhang and colleagues, who showed that AQP4 deletion (homozygous knockout) intensified αSyn pathology and motor impairments as a result of viral-vector delivered A53T-αSyn into the substantia nigra [31]. Both of these studies suggest a clear role of AQP4 in regulating αSyn accumulation in the brain but are limited by their use of genetic approaches to manipulate glymphatic function *in vivo*; reduced AQP4 expression from birth, present at the time of pathology induction. We show here that (pharmacological) glymphatic inhibition results in a ∼75% greater retention of intracerebrally delivered agents. Since both Cui and colleagues, and Zhang and colleagues’ models were initiated by delivery of substances to the brain, it is quite possible that the exacerbated pathology that both groups observed as a result of AQP4 deletion was in fact the result of enhanced delivery of the initiating agent (AAV or PFFs). In other words, by taking such a genetic approach, it is impossible to isolate the effects of AQP4 reduction on pathological development from its effects on enhanced pathological induction. In our study, glymphatic function was pharmacologically inhibited *after* model establishment; mice receiving TGN-020 treatment starting ∼3 days post-injection with PFFs. Hence, we can infer that the exacerbated pathological progression we saw in mice was a result of enhanced propagation and pathological development of αSyn pathology in the brain, rather than increased seeding, substantiating the suggestion that αSyn propagation and accumulation are regulated by the glymphatic system.

For many years, functional studies of the glymphatic system in relation to neurodegenerative disease pathology were focussed almost exclusively on the clearance of β-amyloid from the brain [e.g. 25, 26], perhaps due to the extracellular nature of β-amyloid plaque pathology. But more recently, studies now implicate failure of the system in the accumulation of a number of intracellular proteins prone to brain-wide propagation [27–31, 58, 59]. For example, we recently published findings on the effects of glymphatic inhibition on tau propagation in a seeded mouse model [51]; enhanced propagation and accumulation of tau in the brain following chronic treatment with TGN-020. In that study, at end-stage following chronic TGN-020 treatment, we saw near off identical levels of exacerbated tau pathology in the hippocampus, to that of αSyn pathology we see here in the striatum, despite different durations of animal studies, different propagation models, and different pathology inducing injectates. Indeed, tau and αSyn are very different proteins (∼55-62 kDa and 14 kDa, respectively), but it appears that their glymphatic clearance profiles are remarkably similar. Disease-associated glymphatic dysfunction and lack of clearance discrimination between neurodegenerative disease associated proteins might explain the mixed or concomitant pathologies which are increasingly reported and characterised in the literature; thought by many to be the norm rather than the exception in late-onset neurodegenerative disease [60–63]. Given the number of protein pathologies the glymphatic system has now been implicated in, studies aiming to hierarchically classify the glymphatic clearance of neurodegenerative disease proteins might provide much needed direction for the field, with the hope of translating potential glymphatic targeting therapies to the clinic.

Aside from the effects of glymphatic inhibition we saw on αSyn pathological development, we also observed regional differences in PFF-induced alterations of glymphatic function, in PFF seeded mice. Glymphatic function in the striatum, which was subject to direct injection of PFFs, was relatively unaffected, but exhibited substantial AQP4 and DAC protein downregulation. Glymphatic function in the midbrain on the other hand, appeared to be upregulated upon receipt of αSyn, with AQP4 expression alterations found only at end-stage in the model. By contrast, Zhang et al. saw reduced glymphatic tracer influx in the whole brain 4 weeks after AAV-A53T injection into the substantia nigra, and reduced AQP4 polarisation in both the striatum and substantia nigra [31]. Similarly, in aged A53T transgenic mice, Zou and colleagues also saw reduced glymphatic inflow in the midbrain, but interestingly also observed increased AQP4 expression, but reduced polarisation in this region [64]. The major difference between both of these studies and ours is the method by which αSyn pathology is induced in the brain. Both Zhang and colleagues, and Zou and colleagues use a genetic approach to drive protein aggregation (via AAV injection or transgenic expression, respectively). Whereas, here, we induced pathological accumulation and propagation of αSyn via PFF injection. Although we perform these experiments in A53T mice, pathological seeding is induced in young animals, where transgene driven αSyn deposition is yet to occur [49]; transgenic expression of pre-pathological aggregate-prone αSyn accelerating rather than initiating pathology in the brain. The discrepancy between our results and both of the aforementioned studies suggests that propagating αSyn, and genetically overexpressed mutant αSyn, have marked differences in terms of their effects on glymphatic function. For example, in the current study we saw that when αSyn pathology physiologically propagates to the midbrain, in contrast to when it is genetically overexpressed and already deposited there, the glymphatic system is able to respond and increase its function. This gives us some insight into what might occur in the Parkinson’s disease brain as αSyn pathology propagates and the disease progressed through the Braak stages [65]; the glymphatic system is plastic in its nature and is able to upregulate its function in response to receipt of aggregate-prone proteins (what we saw in the midbrain in our study). Once protein deposition is established however (like in the striatum in our study, and end-stage in the midbrain) and neurodegeneration is evident, AQP4 expression becomes affected. We can infer from our drug study that reduced AQP4 expression likely exacerbates αSyn pathology, but much is still left to learn about precisely how αSyn, particularly propagating αSyn, affects AQP4 expression and channel function. Understanding the mechanisms by which these two proteins affect one another may provide information as to much needed novel drug targets for potential disease alteration.

Glymphatic function has similarly found to be impaired in non-αSyn based mouse models of Parkinson’s. For example Si and colleagues found glymphatic inflow and tracer clearance to be impaired when dopaminergic neurodegeneration was induced by 1-Methyl-4-phenyl-1,2,3,6-tetrahydropyridine (MPTP) in mice [66]. Interestingly, in this setting, reduced glymphatic function was only seen to be impaired in the midbrain and substantia nigra (the location of the dopaminergic cell bodies undergoing degeneration) but not in the striatum (location of the dopaminergic terminals projecting from the nigra), despite significant declines in tyrosine hydroxylase positive signal observed at both locations. Similarly, in LRRK2 (mutations of which are the most common genetic cause of Parkinson’s, and are associated with high risk of sporadic Parkinson’s disease development [67]) mutant (R1441G) mice, glymphatic function and AQP4 polarisation were observed to be reduced [68]. The authors of this study also show that LRRK2 itself interacts with AQP4 to phosphorylate it, which leads to reduced polarisation of the channel. These studies add further insight into the wealth of potential influences biological perturbations in the Parkinson’s brain may have on the function of the glymphatic system and highlight that although αSyn accumulation plays a substantial role in disease development and progression, its aggregation is not the only factor which likely affects the capacity of the glymphatic system to rid the brain of interstitial waste solutes.

Sold as an AQP4 inhibitor, TGN-020 has now been shown by numerous independent groups to inhibit glymphatic function *in vivo* [66, 69–74]. Indeed, our own group has shown that both acute and chronic treatment with the agent reduces interstitial inflow of CSF tracers (both fluorescent and gadolinium-based) [27, 51]. In xenopus oocytes, TGN-020 has been shown to be isoform specific for AQP4 compared to other aquaporins, with an IC_50_ of 3.1 µM [75]. However reports suggest difficulty reproducing these cell-based assays [76], with lack of drug effect observed in primary astrocytes expressing endogenous AQP4 or mammalian cell-lines overexpressing exogenous AQP4 [77]. Indeed, published *in vivo* studies using TGN-020 fail to clarify its mechanism of action, hence careful interpretation of results is required. In our hands, like many groups, we see robust glymphatic inhibition with TGN-020 *in vivo* [27, 51], but this raises the suggestion as to what other targets TGN-020 is interacting with to affect glymphatic function. AQP4 has been central to studies of the glymphatic system since its discovery, but the clear effect we see here of this drug on αSyn propagation and accumulation underscores the need for more in-depth study of its mechanism of action, *in vivo*, which may lead to identification of additional targets able to influence glymphatic function.

In the current study we made use of a transgenic model of αSyn propagation, in which pathology was seeded in the brain with αSyn PFFs, and propagation of αSyn pathology is accelerated by the transgenic expression of A53T mutant human αSyn. First described in 2012 [54], this model provides a robust platform on which to study mechanisms associated with the propagation of human αSyn in the brain, within a feasible experimental period, as we have here. Achievement of the model is reliant upon generation of suitable αSyn PFF seeding material, which is highly dependent on buffer conditions, sonication efficiency, and storage, hence large batch-to-batch and lab-to-lab variability has been reported, leading to reproducibility issues [44, 78]. To minimise these in our study, we made use of the Michael J. Fox Foundation for Parkinson’s Research (MJFF) guidelines and best practices for generating PFFs [44], ensuring their thorough validation ahead of *in vivo* work and use of a single PFF batch for all work presented. Although within our own study we saw highly reproducible development of αSyn pathology in the brain, we observed substantial discrepancies in terms of the time course of disease phenotype observed in animals injected with our PFFs compared to other groups. Injection of our generated PFFs (5 µg, unilaterally) into the striatum and overlying cortex of M83 mice led to a median survival time (time to reach humane endpoint) of 61 days (data not shown), which was highly uniform as previously noted by others [54]. Hence in the current study we chose to treat animals with the glymphatic inhibitor for 6 weeks (42 days), to allow us to assess the extent to which drug treatment exacerbated pathology in the brain. The period of survival post-PFF injection in our characterisation of the model, was far shorter than Luk and colleagues, who originally described the model. Researchers reported a median survival time of 101 days, despite use of an identical PFF dose, injection site and transgenic mouse line for investigations [54]. The authors of this work also struggled to quantify motor impairments in their mice due to the rapidity of which mice develop symptoms at end stage. Indeed, in vehicle treated PFF seeded mice in our study, we were unable to detect a motor impairment with open field at 6 weeks; only observing apparent locomotor reduction in drug treated animals at this timepoint. Our findings are more in line with Earls et al., who saw clinical symptoms (unstable gait and hunched posture with extensive hind limb retraction) at 70 days post-injection prior to brain collection at the end of their study, with a number of mouse deaths observed prior to this timepoint with their treatment, which exacerbated pathology [79]. Authors of this study interestingly also observed increased limb clasping behaviour in PFF injected M83 mice from 5 weeks post-injection and onwards. The notable discrepancies between studies using this model highlights how important thorough validation of the PFF seeding material is, and shows that although robust in-lab, this model is exceptionally prone to between-lab variability, serving as a warning to researchers to establish this model with caution, and ensure thorough characterisation of it in their own hands before embarking on large experimental cohort studies.

In conclusion, we have shown here that in a mouse model of αSyn propagation, the expression profiles of AQP4 and its endfoot binding partners are affected by phosphorylated αSyn load. Incoming propagation of αSyn in the midbrain however encourages an enhancement of glymphatic function in response, to manage αSyn burden, ahead of AQP4 and endfoot protein dysregulation. Experimental inhibition of glymphatic function in mice reduces brain to CSF clearance of αSyn acutely and exaggerates pathology and behavioural impairments in an extended treatment paradigm. Together our findings suggest that αSyn-induced dysregulation of AQP4 and the DAC at the astrocytic endfoot likely contributes to the glymphatic dysfunction observed in Parkinson’s disease, and that this impairment may exacerbate αSyn accumulation and pathology, establishing a positive feedback loop.

## Data availability

Data available on request.

## Acknowledgements

Authors would like to thank Dr Phillip Muza and Miss Barbara Lechnicka for their critical appraisal of the manuscript.

