## Supplementary material for "Influence of the Glymphatic System on α-Synuclein Propagation: Role of Aquaporin-4 and the Dystrophin-Associated Protein Complex"

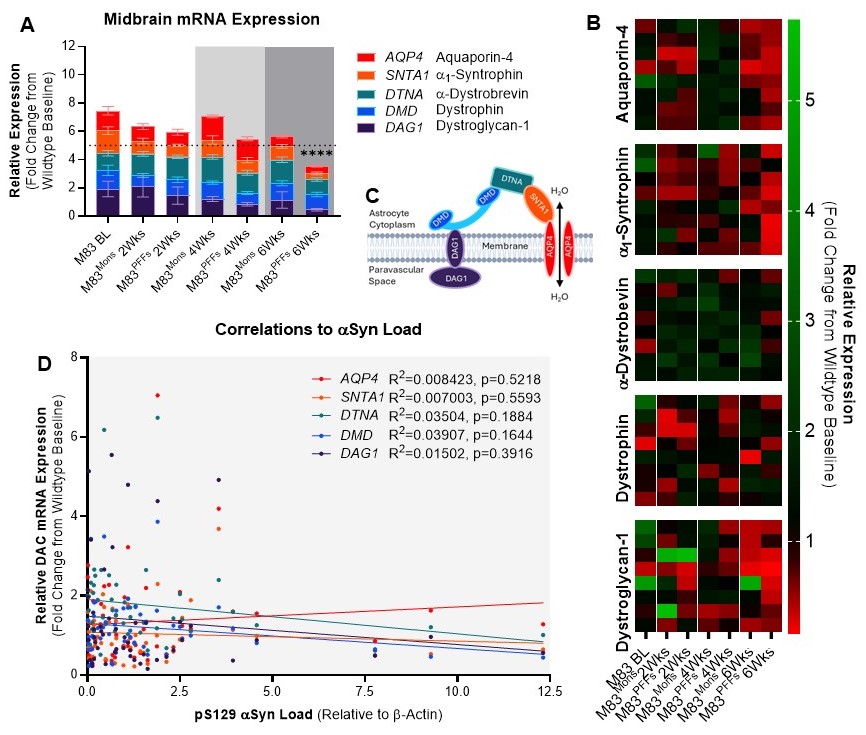
**Supplementary Figure 1 – Propagation of αSyn pathology to the midbrain induces dystrophin associated complex dysregulation at only 6 weeks following PFF injection, with no overall correlation to αSyn load**

(**A**) Midbrain mRNA expression levels of AQP4 and DAC elements in experimental groups injected with either αSyn monomers or PFFs and left to age for 2, 4 or 6 weeks. Expression data is normalised to wildtype baseline expression levels (dotted line). Individual animal datapoints (n=8) shown in heatmap (**B**). (**C**) diagrammatic illustration of the DAC at the astrocytic endfoot membrane, with elements colour-coded to reflect data in (A). Correlation analysis of mRNA expression profiles with animal-by-animal αSyn load data, illustrating lack of any discernible correlations between AQP4 DAC element expression and αSyn load. Results from statistical tests (B, 2-way ANOVA; D, simple linear regression fitting) indicated with asterisks: ****=p<0.0001. Asterisks in (A) indicate significance from M83 baseline (BL).

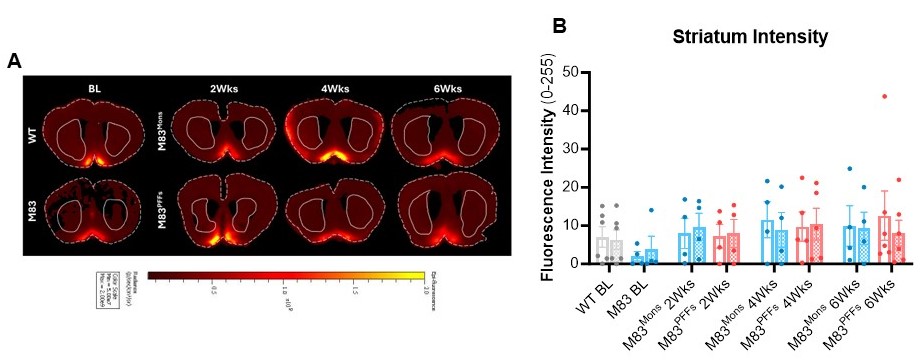
**Supplementary Figure 2 - αSyn PFF seeding does not result in altered glymphatic function in the striatum**

(**A**) Fluorescence intensity in the striatum (solid outline) in brain sections (dashed outline) following cisterna magna infusion of Texas Red™-conjugated dextran (3 kDa). Intensity is quantified and graphed in (**B**) illustrating lack of any discernible differences between groups. The ipsilateral striatum area is illustrated with empty bars, the contralateral striatum area is illustrated with hashed bars. Statistical tests (B, 2-way ANOVA) indicated no significant differences.

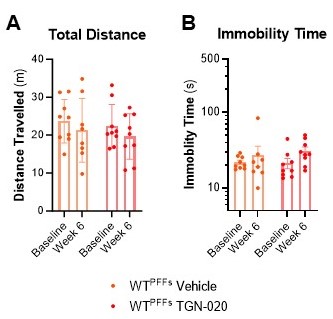

**Supplementary Figure 3 – TGN-020 does not elicit motor impairments in wildtype mice following PFF injection**

No behavioural impairments are observed in wildtype (WT) mice injected with PFFs. And addition of TGN-020 treatment does not elicit a behavioural response in either total distance (**A**) or immobility time (**B**) in the open field arena. Statistical tests (2-way ANOVAs) indicated no significant differences. Data is graphed on the same y-axis scale as data presented in Fig. 5C and D, for direct comparison between datasets.
